# centriflaken: an automated data analysis pipeline for assembly and *in silico* analyses of foodborne pathogens from metagenomic samples

**DOI:** 10.1101/2025.07.18.665485

**Authors:** Kranti Konganti, Julie Kase, Narjol Gonzalez-Escalona

**Author notes:** Corresponding author. Mailing address, Human Foods, Program, Food and Drug Administration, 5001 Campus Drive, College Park, MD 20740, USA.

## Abstract

Rapid and comprehensive analysis of metagenomic data from samples associated with foodborne outbreaks is of critical importance in food safety. Equally important is the need for automated analysis pipelines that allow the rapid and effective construction of metagenomic assembled genomes (MAGs) to enable bacterial source-tracking from metagenomic data. Here, we present centriflaken, an automated precision metagenomics pipeline for detecting and characterizing Shiga toxin-producing *Escherichia coli* (STEC) from metagenomic data. centriflaken streamlines the process of generating metagenome-assembled genomes (MAGs) and conducting *in silico* analyses of STECs, significantly reducing the time and manual effort required for comprehensive pathogen profiling. centriflaken was validated using Oxford Nanopore long-read sequencing data from agricultural water enrichments, successfully reproducing results from our previous study that involved multiple manual bioinformatics steps (Maguire et al., 2021). The tool’s efficacy was further demonstrated through its application to ZymoBIOMICS microbial community standards and 21 additional irrigation water samples, completing STEC precision metagenomics analyses in less than 7 hours per sample. centriflaken’s versatility allows for the analysis of user-defined taxa beyond STEC, including other foodborne pathogens like *Listeria monocytogenes* or *Salmonella*. The pipeline generates comprehensive summary plots and tables, accessible through a MultiQC HTML report. Designed for portability, centriflaken packages all software dependencies within containers and virtual environments. This open-source tool is available on GitHub under the MIT license (https://github.com/CFSAN-Biostatistics/centriflaken), offering a powerful resource for rapid, automated pathogen detection and characterization in food safety applications.

**Author summary:** Metagenomic sequencing, particularly using nanopore technology, generates vast amounts of data that are challenging to analyze, especially when searching for specific pathogens like Shiga toxin-producing *Escherichia coli* (STEC). This challenge is compounded when processing multiple samples simultaneously. The pipeline “centriflaken” was developed to streamline this complex process, automating the extraction of *E. coli* reads from metagenomic data and performing *in-silico* characterization of potential STEC present in the samples. centriflaken builds on the concept of precision metagenomics, employing a suite of automated data analysis workflows powered by Nextflow. The pipeline processes metagenomic data, generates metagenome-assembled genomes (MAGs), and conducts *in-silico* analyses as described in Maguire et al. (2021). By running steps in parallel, centriflaken enables users without extensive bioinformatics skills or STEC genomic knowledge to efficiently analyze complex metagenomic data. Key features of centriflaken include: 1) Automated workflow for STEC detection and characterization, 2) Parallel processing for improved efficiency, 3) User-friendly interface for non-specialists, 4) Scalability for analyzing multiple samples simultaneously, and 5) Compatibility with high-performance computing (HPC) clusters and cloud environments. This freely available software democratizes complex metagenomic analysis, making it accessible to a broader range of researchers and food safety professionals.

## INTRODUCTION

As farm-to-table food trends increase in popularity, we have seen an increase in produce related outbreaks of foodborne illness. Shiga toxin-producing *E. coli* (STEC) are a major source of foodborne illness [1–4]. In fact, O157:H7 accounts for 70% of *E. coli* related foodborne outbreaks [5–10]. Recent outbreaks underscore the ongoing threat of STEC contamination. For instance, in October 2024, a multi-state outbreak caused by a O121:H19 STEC strain linked to organic carrots resulted in 48 illnesses, 20 hospitalizations, and 1 death, highlighting the critical need for rapid and accurate STEC detection and characterization methods (https://www.cdc.gov/ecoli/outbreaks/e-coli-o121.html).

Due to the ubiquitous nature and the varying degrees of pathogenicity of *E. coli*, its mere presence does not equate to concerns to public health, unlike with bacteria such as *Salmonella* or *Listeria monocytogenes*. Therefore, the detection of *E. coli* presence alone is insufficient to assess health risk. Moreover, there are over 400 described STEC serotypes that express one or more of *stx*1 genes and *stx*2 genes [11, 12] and among those, around 100 serotypes have been described as causing illness in humans, with hemolytic uremic syndrome (HUS) being deadly [13–18]. However, serotype and *stx* gene presence and variant type are not enough to predict STEC risk for humans. STEC must also have other virulence genes linked to attachment and colonization of the intestine such as intimin (*eae* gene), AggR (*aggR* gene), auto-agglutinating adhesin Saa (*saa* gene), chromosome located T3SS effectors encoded by genes such as *tir, espA, espB, espK,* among others, and other virulence genes located in plasmids (*toxB, hexA, etpD, katP*) [19–23].

In our previous study, we established the use of nanopore sequencing to determine the presence and full characterization of STECs in enrichments of agricultural water [24]. We determined that the limit for obtaining a complete, fragmented assembly for STECs by nanopore sequencing of DNA extracted from enriched agricultural water was 10^5^ CFU/ml [24]. There are many advantages of using nanopore sequencing for STEC identification and characterization: affordability, potential to multiplex, portability, less manual labor required from personnel, use of low DNA concentration, longer reads than other available technologies (>100 kb) and the analysis could also be done in real-time (using their metagenomics workflow available at EPI2ME labs).

A previous pilot study produced preliminary evidence that MinION sequencing of agricultural water using the ligation kit has the potential to be used for rapid microbiome determination in the field with optimal results for water quality surveillance [25]. Agricultural water has been implicated as a source of contamination for produce resulting in foodborne illness and outbreaks [26–31]. Current FDA protocols for the detection and isolation of STECs require multiple rounds of selective plating and whole genome sequencing (WGS) of a single isolate. On-site field testing is increasingly becoming a priority to decrease the time for detection of pathogenic microbes and for the fast implementation of corrective measures. While the most abundant species in the microbiome is likely to fluctuate seasonally, they may be an indicator to changing populations and importantly may serve to monitor deviations in microbial water quality.

However, any of these studies or experiments will generate a large amount of data. Analyzing nanopore data or any other high-throughput data is a very complex procedure and usually requires many steps to generate the final reportable results creating a bottleneck. Metagenomic workflows are usually compute intense and need to be analyzed on High-Performance Compute Clusters (HPCs). In the case of data generated from metagenomic samples, tools are needed to detect the presence of STEC and comprehensively assess their virulence potential bysubtyping the *stx* genes, determining serotype, identifying attachment genes such as *eae* and *saa* and detecting other virulence-related genes. A precision metagenomics approach is not meant to replace the cultural method [32], but it is a way to detect and identify a STEC strain earlier and evaluate its potential risk to the public. Improving STEC genome characterization methods has significant implications for public health and food safety. Rapid, accurate identification and genome characterization of STECs can lead to faster response times in outbreak situations, more targeted recalls, and ultimately, fewer cases of foodborne illness. Moreover, enhanced detection capabilities could inform more precise and effective preventive controls in food production and processing, aligning with FDA’s risk-based approach to food safety.

The existing protocol or pipeline in our FDA laboratory uses many sequential manual steps, described in Maguire et al. (2021) [24]. Briefly, the steps are divided into: 1) classify reads using WIMP workflow available at EPI2me, 2) a python script to extract or bin reads that matched the requested taxa (i.e. *E. coli*), 3) Assemble the extracted reads using flye assembler [33], 4) Classify the contigs with Kraken 2 [34] and extract contigs matching requested taxa, 5) perform all *in silico* analysis on those contigs (MLST, AMR, serotyping, and virulotyping), and 6) aggregate all data into a final report and conduct a risk assessment for the sample.

Several other workflows for metagenomic analyzes have been published and reviewed by Van Damme et al. 2021 [35]. These authors described the creation of a very useful metagenomic tool that allows for the analyses of metagenomic data using both short and long reads in a hybrid approach which allows for the reconstruction of more complete and more accurate metagenome-assembled genomes (MAGs) [35]. While these approaches have strengths, none of them allows for a precision metagenomic analysis (taxa specific) and *in silico* virulotyping, serotype, and AMR prediction for STECs in a single workflow.

Here we present centriflaken to address this gap. centriflaken is an integrated suite of automated data analysis workflows enabled by Nextflow [36]. This pipeline processes metagenomic data to generate metagenome-assembled genomes (MAGs) and performs comprehensive *in silico* analysis for Shiga toxin-producing *Escherichia coli* (STEC), following the approach described by Maguire et al. (2021). centriflaken streamlines the entire process, from raw sequencing data to final results, and provides user-friendly output in the form of summary plots and tables, accessible through a MultiQC HTML report generated automatically by the pipeline. The development of centriflaken aligns closely with FDA’s New Era of Smarter Food Safety initiative, which emphasizes the use of new technologies and approaches to create a safer, more digital, and traceable food system. By enabling faster, more comprehensive analysis of metagenomic data, centriflaken supports FDA’s goals of strengthening root cause analyses, improving predictive analytics, and responding more rapidly to outbreaks.

## MATERIALS AND METHODS

### Centriflaken pipeline

A high-level overview of centriflaken workflow is presented in Figure 1. The pipeline can be run on any UNIX based machine. The main analysis starts by taking UNIX input paths to the fastq_pass reads folder generated by Oxford Nanopore. It then filters out any reads less than 4000 bp and then performs taxonomic classification and extracts all *E. coli* only reads using centrifuge [37] followed by taxon specific assembly using flye assembler v2.9 [33]. The assembled contigs are reclassified to identify the *E. coli* taxon using kraken2 [34] and extract the *E. coli* contigs to produce a final Metagenomically assembled Genome (MAG). The final MAG is interrogated (in the case of STEC) against several specific databases.

**Figure 1.**
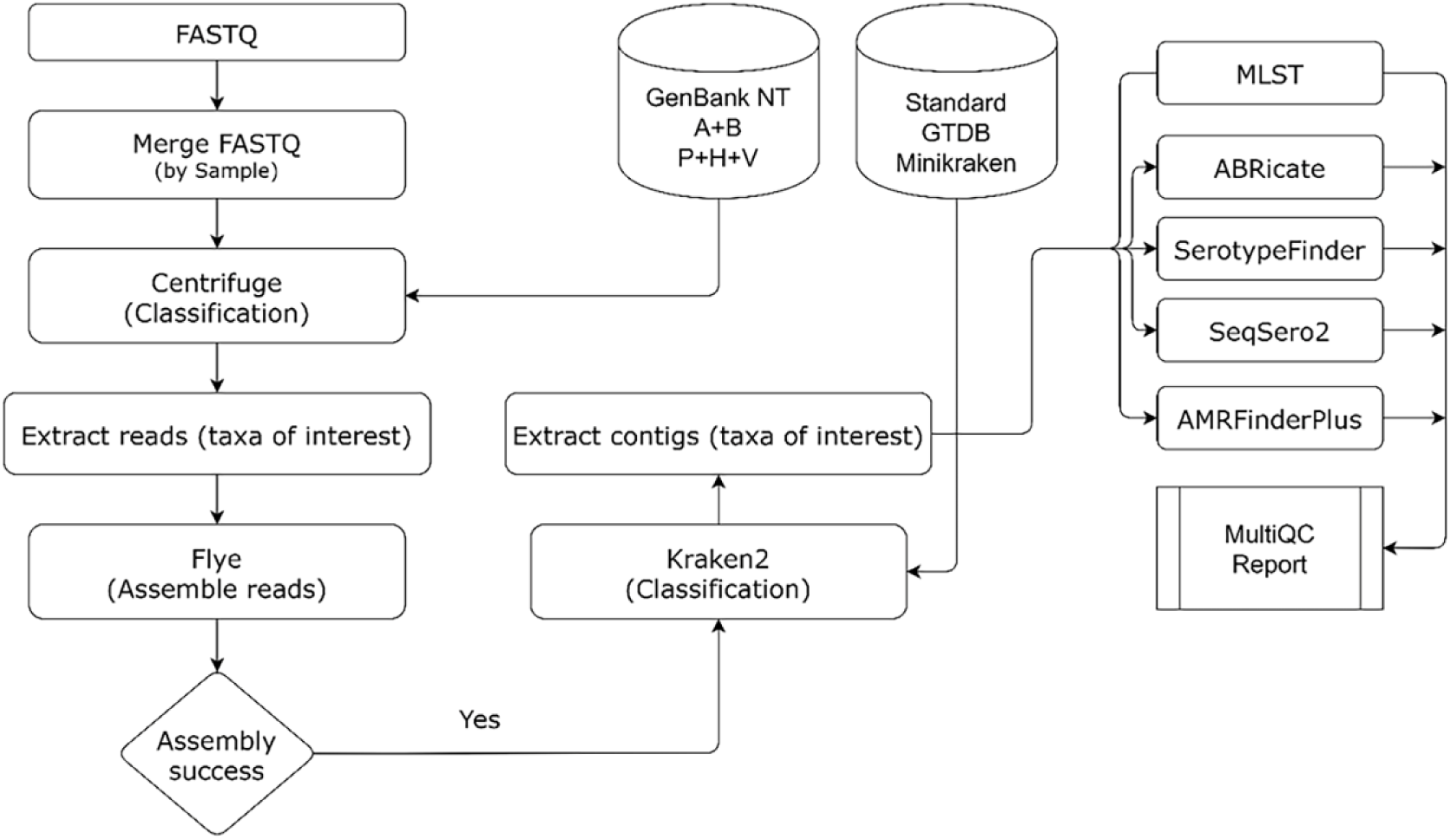
A brief overview of the centriflaken pipeline. The pipeline starts by taking as input a UNIX path to FASTQ files after which multiple read files belonging to the same sample are merged followed by excluding reads whose length is less than 4000 bp. Then, reads are extracted belonging to a taxon of interest using Centrifuge following which the target assembly is performed using flye. Kraken2 is now used to extract contigs binned as the taxa of interest and these contigs are used for further downstream analysis such as subtyping, virulotyping, AMR gene finding, etc. The outputs from all these downstream processes are used to generate a summary report using MultiQC.

Serotypefinder [14] is used to serotype the MAGs, ABRicate (https://github.com/tseemann/abricate) is run to identify known AMR genes using various databases (AMRfinder plus [38], MEGARes 2.0 [39], RESFINDER [40], ARG-ANNOT [41]) and their results are shown in a final MultiQC report. All the analysis steps are automated using Nextflow and the pipeline runs in parallel for all samples. The pipeline took approximately 6 hours to finish processing the data produced in Maguire et al. (2021) [24], which previously took many days to run all steps for all samples manually or through automated bash scripts.

The Centriflaken GitHub repository provides guidance on acquiring the necessary Centrifuge and Kraken databases. For Centrifuge, links to pre-built indices are available, while for Kraken, step-by-step instructions for database construction are provided. Detailed information can be found in the repository’s README file at: https://github.com/CFSAN-Biostatistics/centriflaken/blob/main/README.md#databases. It is recommended to run centriflaken using conda software virtual environments for users who do not have access to HPC cluster with singularity. The pipeline can also be run using docker or singularity containers by switch the -profile flag when invoking the pipeline. Detailed instructions on setup and running the pipeline on data from Maguire et al. 2021 [24] can be found at https://github.com/CFSAN-Biostatistics/centriflaken/blob/main/readme/centriflaken.md. centriflaken is also made available via Galaxy user interface via the FDA HFP GalaxyTrakr instance (https://galaxytrakr.org). The previously published protocol (https://www.protocols.io/view/centriflaken-an-automated-data-analysis-pipeline-f-kxygxzdbwv8j/v5) can be used to run analysis on GalaxyTrakr. While Centriflaken is designed to work with both Illumina short reads and Oxford Nanopore long reads, this publication specifically focuses on its application to long-read sequencing data generated by Oxford Nanopore technologies.

### Validation of centriflaken and accuracy assessment

Centriflaken was initially tested and validated using data from Maguire et. al. 2021 [24]. The presence of the complete genome and synteny of the completely closed genomes on the final assemblies was checked using the whole genome alignment tool available in the CLC Genomics workbench software v24.0 (QIAGEN, Redwood City, USA). Further testing was conducted with data generated from two ZymoBIOMICS Microbial Community Standards (ZymoBIOMICS Microbial Community Standard II - log distribution - catalog number D6311, and ZymoBIOMICS Microbial Community DNA Standard - catalog number D6306) (ZymoBIOMICS, Irvine, CA), and can be found on GenBank using the accession numbers in supplementary Table 1. The centriflaken final testing was done using 21 overnight enriched samples for STEC collected in two different seasons during 2020 and 2021.

### Overnight enrichments of agricultural waters

Water samples were collected and prepared per this protocol: dx.doi.org/10.17504/protocols.io.j8nlk8nydl5r/v1. Briefly, water samples (225 ml) from the Southwestern US were enriched by adding an equal volume of 2× modified Buffered Peptone water with pyruvate (mBPWp; NEOGEN, Lansing, MI, USA) as described in USFDA Bacteriological Analytical Manual (BAM) Chapter 4A [32], with 6.0 X mL of acriflavin-cefsulodin-vancomycin (ACV) supplement added after five hours of static incubation at 37°C ± 1°C. After the addition of the ACV supplement, enrichments were incubated at 42°C ± 1°C for an additional 16 ±3 hours.

### DNA extraction

Genomic DNA from 1ml of each of the 21 enriched agricultural water sample was extracted using the Maxwell RSC Cultured Cell DNA Kit with a Maxwell RSC Instrument (Promega Corporation, Madison, WI, USA) according to the manufacturer’s instructions DNA concentration was determined by a Qubit 4 Fluorometer (Invitrogen, Carlsbad, CA, USA) according to the manufacturer’s instructions.

### STEC qPCR detection

The presence of STEC was determined by qPCR as described in Chapter 4A of the FDA BAM detecting *stx1*, *stx2*, and *wzy* [32]. Briefly, the DNA retrieved from the 21 samples were diluted 1:10 in nuclease-free water and 2µl was added to 28µl master mix containing 0.25µM stx1 and stx2 primers, 0.3µM wzy primers, 0.2µM stx1 probe, 0.15µM stx2 and wzy probes, 1X Internal Positive Control Mix (Cat: 4308323, Applied Biosystems), 1X Express qPCR Supermix Universal Taq (Cat: 11785200, Invitrogen), and ROX passive dye. All primers and probes (Supplementary Table 2) employed in this study were purchased from IDT (Coralville, IA, USA).

### Nanopore sequencing

DNA recovered from the 21 enriched AW samples was sequenced using a GridION nanopore sequencer (Oxford Nanopore Technologies, Oxford, UK). The sequencing libraries were prepared using the Genomic DNA by Ligation kit (SQK-LSK109) and run in FLO-MIN106 (R9.4.1) flow cells, according to the manufacturer’s instructions for 72 hours (Oxford Nanopore Technologies). The runs were live base called using either Guppy v3.2.10 or 4.2.3 included in the MinKNOW v19.12.6 and v20.10.6 software (Oxford Nanopore Technologies, Oxford, UK), respectively, with the fast-calling model. The two ZymoBIOMICS Microbial Community Standards (ZymoBIOMICS Microbial Community Standard II - log distribution - catalog number D6311, and ZymoBIOMICS Microbial Community DNA Standard - catalog number D6306) (ZymoBIOMICS, Irvine, CA), were also sequenced as described above using 1 ug of DNA as the initial amount for preparing the library.

### Data availability

The raw ONT data for the 7 samples used for the initial validation can be found under Bioproject PRJNA639799. The raw ONT data generated for each individual sample in this study (ZymoBIOMICS samples and the 21 AW) were deposited in GenBank under BioProject accession number PRJNA1222809. Each sample SRA number matching its corresponding sample can be found in Supplementary Table 1.

## Results

### Software Pipeline Overview

The data analysis steps in centriflaken are automated using the Nextflow workflow manager [36], which provides advantages such as process parallelization, sample provenance, retry on failure, tool dependency solutions using containers and conda software virtual environments among many others. Successful execution of the workflow produces output for each process in its own output folder named after the process name. The workflow design also ensures reproducibility of results, and it is easy to share HTML brief reports for further assessment. Due to the inherent advantages of using Nextflow, centriflaken can be run on-premises HPC clusters or in the cloud.

### Initial Validation with Known samples

The entire centriflaken workflow was initially executed and validated on 7 samples from the Bioproject PRJNA639799 (available at NCBI) using conda software virtual environments. This metagenomic study provides nanopore-based data for each water enriched sample spiked with different STEC levels to determine detection and classification of STECs using nanopore sequencing [24]. These nanopore read dataset provided differential coverage for the spiked STEC strain, and the workflow runtime for all seven samples was around 20 hours. The entire process when performed manually took almost 24 hours for a single sample [24]. centriflaken reduced the analysis time from 24 hours per sample to approximately 3 hours per sample, representing an 87.5% reduction in processing time. centriflaken recovered STEC genomes with genome completeness of at least 85% when the concentration of spiked STEC was above 10^6^ CFU/ml (Table 1 and Figure 2). For comparison, the number of total contigs per STEC spiked water sample when performing the step by step analysis nanopore data workflow [24] was a little bit higher than when using centriflaken, but the same number of contigs containing the STEC spiked genome was recovered, confirming the equivalency of both analysis workflows (Table 1). However, centriflaken has a step that extracts the contigs matching the requested taxa and therefore the analysis is only performed in those contigs, accelerating and making the analysis more precise (see workflow on Figure 1). The pipeline analysis also showed the same virulence profile for each sample to what was reported previously in Maguire et al 2021 (Table 2, supplementary MultiQC HTML report 1), demonstrating the accuracy and reproducibility of the centriflaken workflow. This high level of genome completeness at concentrations above 10^6^ CFU/ml demonstrates centriflaken’s sensitivity and accuracy in detecting and characterizing STEC in complex metagenomic samples, a crucial capability for food safety applications

**Figure 2.**
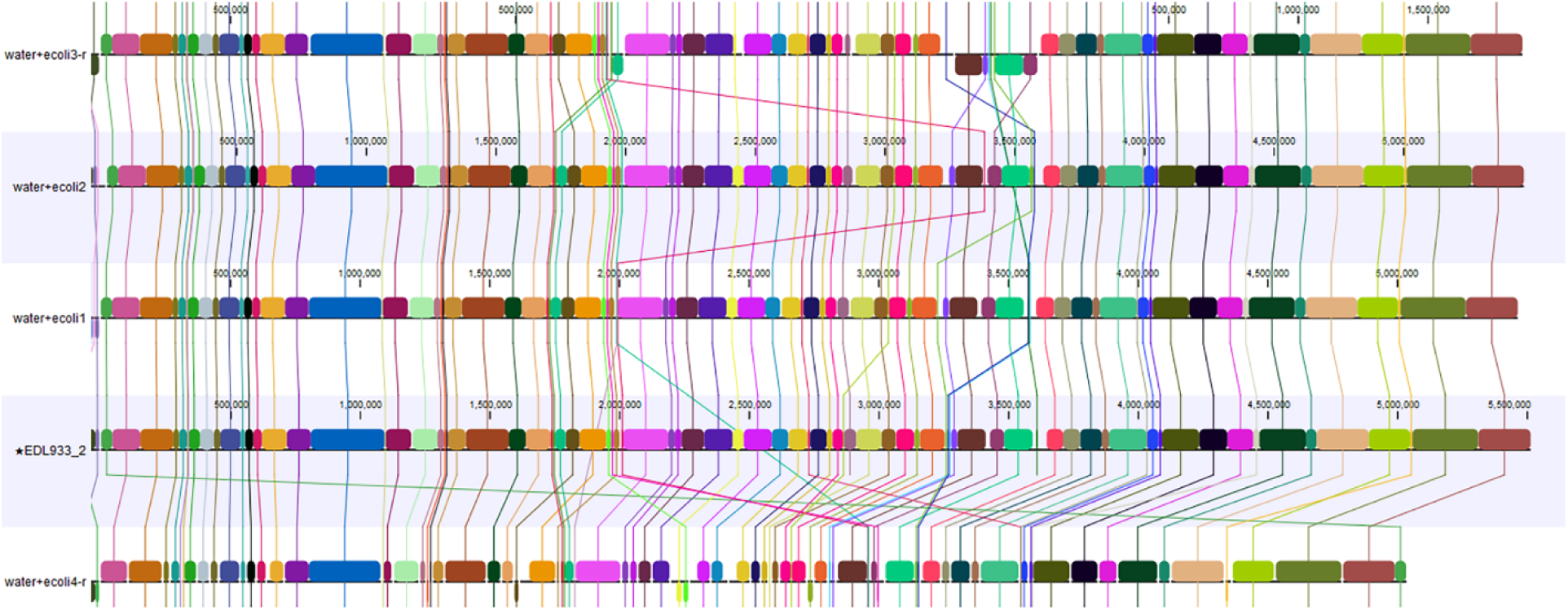
Comparative analysis of EDL933 genome assemblies obtained by centriflaken during the initial validation with known samples. This figure compares the EDL933 genome of the strain used in Maguire et al. (2021) with assemblies obtained using centriflaken for the same samples at various EDL933 enrichment spiking levels. The analysis demonstrates the recovery of the *E. coli* O157:H7 MAG (Metagenome-Assembled Genome) in either completely closed or fragmented forms (see Table 1). Each horizontal track represents EDL933-matching contigs extracted from a sample. Homologous segments across genomes are indicated by the same color and connected. Sequence coordinates in base pairs are shown on respective scales. The reference genome, labeled as EDL933_2 and indicated with a star (*), serves as the basis for comparison.

**Table 1.**
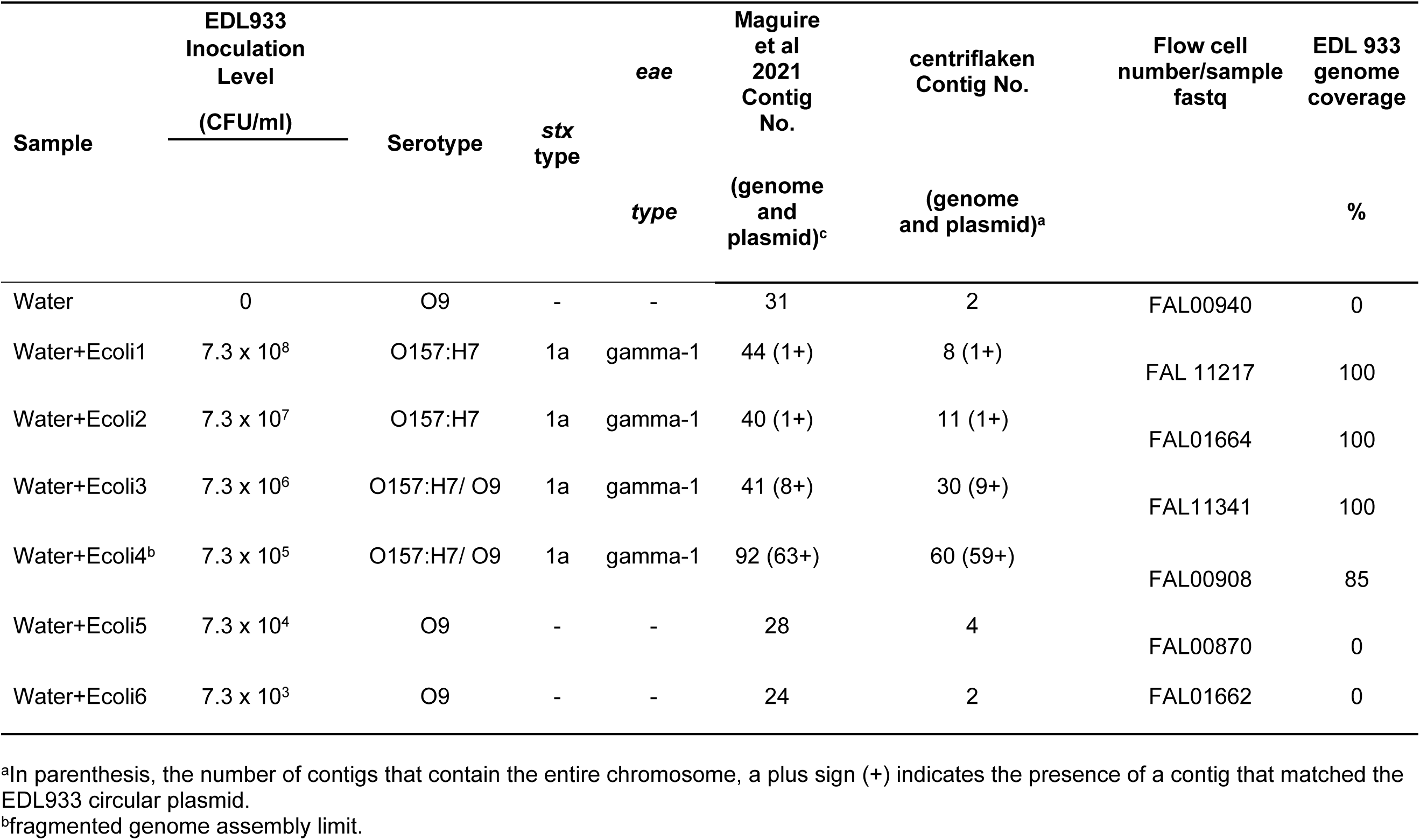
*In silico* virulence factor analysis of metagenomic assembled genomes (MAGs) obtained using centriflaken for the Maguire et al. (2021) dataset. Showing the agreement of the results obtained by both workflows (manually vs. centriflaken)

**Table 2.** Comparative results of the virulotyping of the same ONT data generated (Maguire et al 2021) [24] by both workflows (manually vs. centriflaken)

|  |  | *E. coli* Maguire 2021 |  |  |  |  |  |  | *E. coli* centriflaken |  |  |  |  |  |  |
| --- | --- | --- | --- | --- | --- | --- | --- | --- | --- | --- | --- | --- | --- | --- | --- |
| Virulence Gene | EDL933 reference | Water<sup>a</sup> | 1 | 2 | 3 | 4 | 5 | 6 | Water | 1 | 2 | 3 | 4 | 5 | 6 |
| *astA* | + | - | + | + | + | + | - | - | - | + | + | + | + | - | - |
| *ehxA* | + | - | + | + | + | + | - | - | - | + | + | + | + | - | - |
| *espA* | + | - | + | + | + | + | - | - | - | + | + | + | + | - | - |
| *espB* | + | - | + | + | + | + | - | - | - | + | + | + | + | - | - |
| *espF* | + | - | + | + | + | + | - | - | - | + | + | + | + | - | - |
| *espJ* | + | - | + | + | + | + | - | - | - | + | + | + | + | - | - |
| *espK* | + | - | + | + | + | + | - | - | - | + | + | + | + | - | - |
| *espP* | + | - | + | + | + | + | - | - | - | + | + | + | + | - | - |
| *tccP* | + | - | + | + | + | + | - | - | - | + | + | + | + | - | - |
| *etpD* | + | - | + | + | + | + | - | - | - | + | + | + | + | - | - |
| *gad* | + | - | + | + | + | + | - | - | - | + | + | + | + | - | - |
| *iha* | + | - | + | + | + | + | - | - | - | + | + | + | + | - | - |
| *iss* | + | - | + | + | + | + | - | - | - | + | + | + | + | - | - |
| *nleA* | + | - | + | + | + | + | - | - | - | + | + | + | + | - | - |
| *nleB* | + | - | + | + | + | + | - | - | - | + | + | + | + | - | - |
| *nleC* | + | - | + | + | + | + | - | - | - | + | + | + | + | - | - |
| *tir* | + | - | + | + | + | + | - | - | - | + | + | + | + | - | - |
| *katP* | + | - | + | + | + | + | - | - | - | + | + | + | + | - | - |
| *pssA* | + | - | + | + | + | + | - | - | - | + | + | + | + | - | - |
| *air* | + | - | + | + | + | + | - | - | - | + | + | + | + | - | - |
| *toxB* | + | - | + | + | + | + | - | - | - | + | + | + | + | - | - |

<sup>a</sup>Samples: Water (0 CFU/ml STEC EDL933), 1 ( $7.3 \times 10^8$ CFU/ml), 2 ( $7.3 \times 10^7$ CFU/ml), 3 ( $7.3 \times 10^6$ CFU/ml), 4 ( $7.3 \times 10^5$ CFU/ml), 5 ( $7.3 \times 10^4$ CFU/ml), and 6 ( $7.3 \times 10^3$ CFU/ml).

### Performance with ZymoBIOMICS Microbial Communities

The centriflaken workflow was further tested on samples from a previous study using the ZymoBIOMICS microbial community DNA Bioproject PRJNA751542, Biosample SAMN20520875 [25] and two additional samples (ZymoBIOMICS Microbial Community DNA Standard and ZymoBIOMICS Microbial Community Standard II - log distribution) (NCBI accession provided here). Both communities consist of a mixture of 10 microbes belonging to different genera and species but at different proportions. ZymoBIOMICS microbial community composition is at a 12% concentration each bacterium and 2% for the two yeasts, while ZymoBIOMICS Microbial Community Standard II composition is quite different. ZymoBIOMICS Microbial Community Standard II composition is as follows: of *Listeria monocytogenes* - 89.1%, *Pseudomonas aeruginosa* - 8.9%, *Bacillus subtilis* - 0.89%, *Saccharomyces cerevisiae* - 0.89%, *Escherichia coli* - 0.089%, *Salmonella enterica* - 0.089%, *Lactobacillus fermentum* - 0.0089%, *Enterococcus faecalis* - 0.00089%, *Cryptococcus neoformans* - 0.00089%, and *Staphylococcus aureus* - 0.000089%.

Only the ZymoBIOMICS microbial community DNA samples contained enough *E. coli* (10^8^ CFU per 1 ug) to be within the limits of genome assembly (>10^6^ CFU/ 1 ug) as establish previously for ONT metagenomic samples using the LSK-109 library preparation kit [24] (supplementary MultiQC HTML report 2). centriflaken identified the *E. coli* presence in the sample belonging to serotype O33:H32 and carried 4 known AMR genes using the NCBI AMRfinder plus tool (*blaEC, mdtM, emrD,* and *acrF*) [38]. The ZymoBIOMICS Microbial Community Standard II - log distribution contained ∼ 10^5^ *E. coli* CFU per 1 ug, well below the assembly limit for the technique. That sample still returned a signal for *E. coli* (159 unique taxa matching reads) but no *E. coli* MAGs were recovered and therefore the correct identity as well as the virulence or serotype characteristics of the *E. coli* strain could not be determined (supplementary MultiQC HTML report 2). These results highlight centriflaken’s ability to accurately identify and characterize *E. coli* strains even in diverse microbial communities, while also demonstrating its limitations at lower bacterial concentrations. This information is vital for understanding the tool’s applicability in various real-world scenarios.

### Analysis of Unknown Agricultural Water

After validating the centriflaken workflow with known samples, the next step was to assess its performance with ONT data obtained after sequencing unknown overnight enriched agricultural water samples suspected of containing STECs. For that purpose, we tested the performance on 21 samples obtained after overnight enrichment of agricultural water. The entire precision metagenomics STEC analysis of the 21 samples was completed in less than 7 hours using centriflaken (supplementary MultiQC HTML report 3). The total reads per experiment ranged from 1.3 to 2.8 million reads, of which 11 to 40% matched *E. coli*, with median read sizes between 4,252 to 6,177 bp (Table 3). The number of total *E. coli* contigs per sample ranged from 178 to 1,069.

**Table 3.**
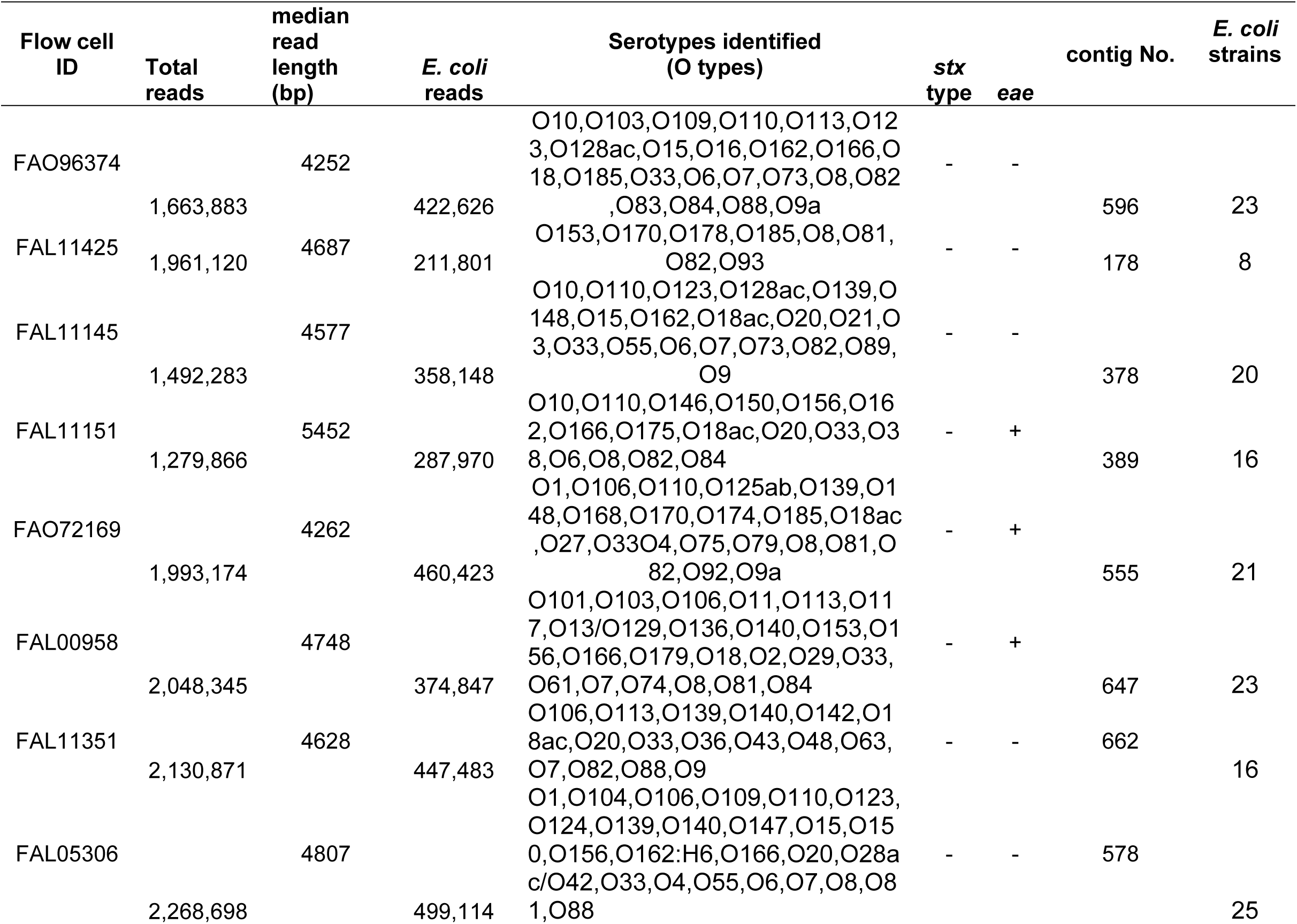
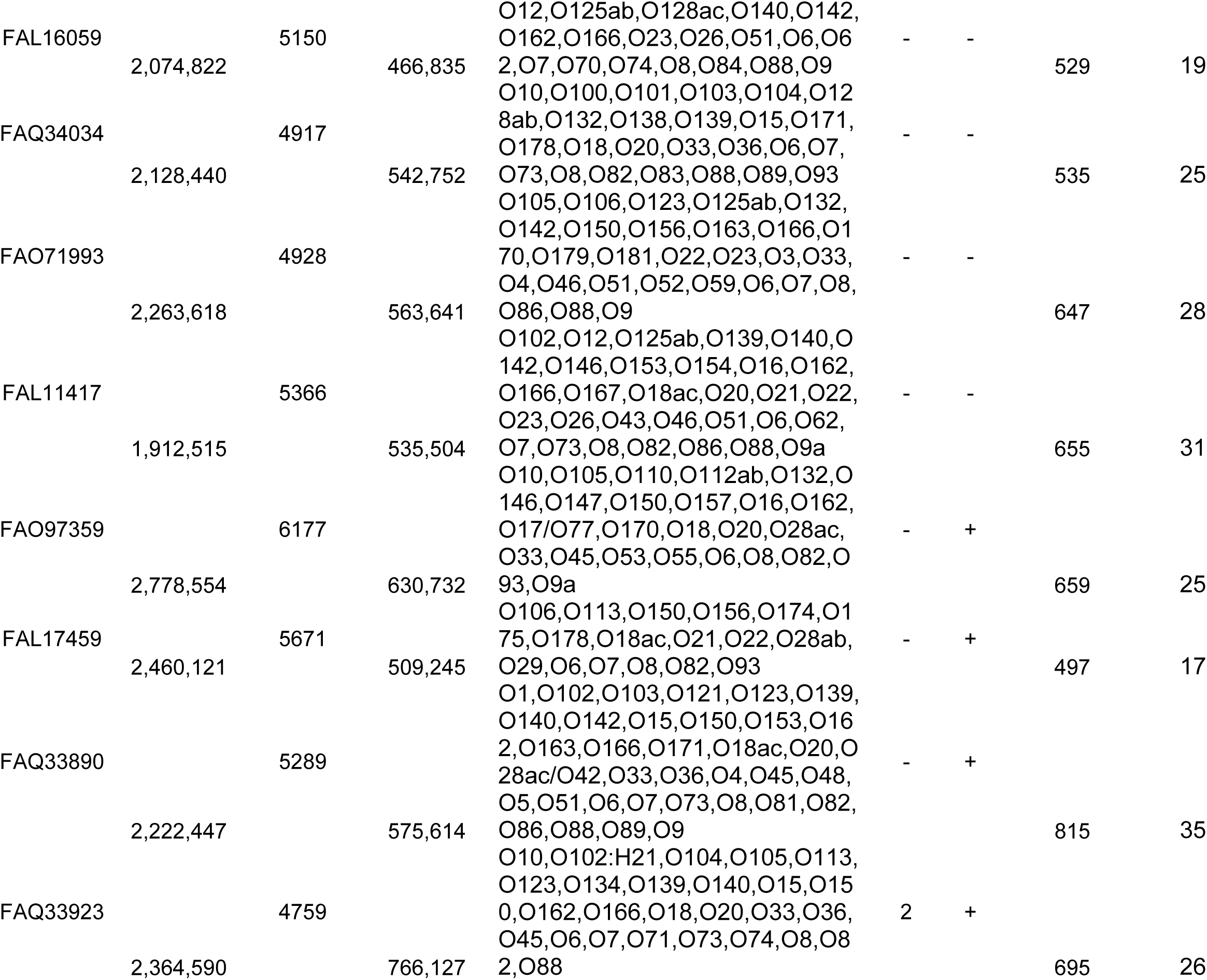
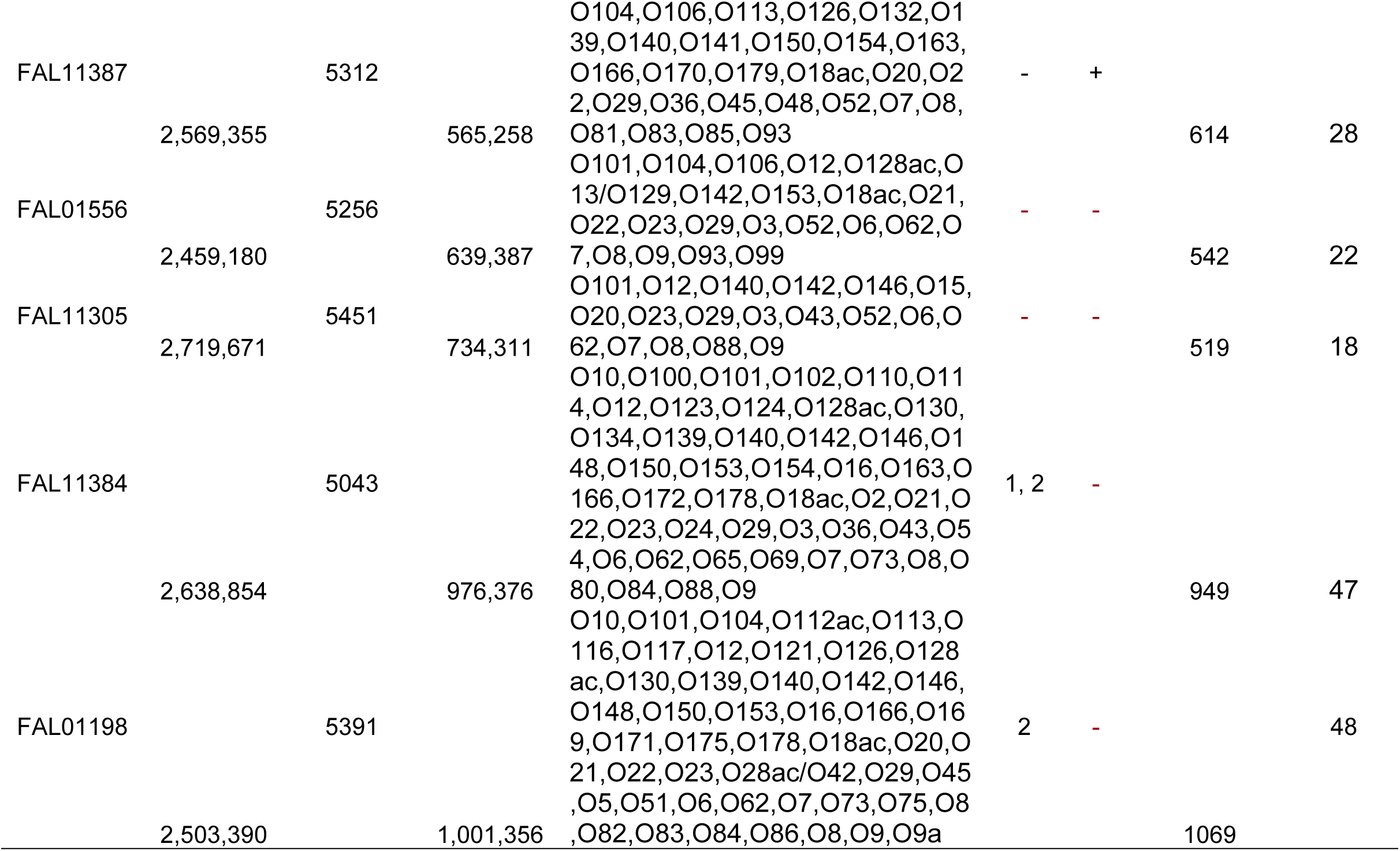
Sequencing statistics, serotypes, *stx* genes and *eae* genes, identified per sample.

Every sample contained more than one different *E. coli* strain, with varying degrees of diversity. If the predicted serotypes identified among the many MAGs obtained per sample is used as a measure of *E. coli* diversity, the number of *E. coli* strains per sample ranged from eight to forty-eight (Table 3). The identification of multiple *E. coli* strains in each sample showcases centriflaken’s capacity to detect and differentiate various strains in complex environmental samples, a key advantage for comprehensive food safety assessments. There were only 3 samples that were identified as positive for the *stx* genes but one of them (FAL11384) lacked any adhesion genes used in our search (Table 3 and 4). Sample FAL01198 was positive for *subA* and FAQ33923 detected the *eae* gene allele epsilon-7. On the other hand, 8 samples returned a signal for the *eae* gene, and one sample contained *saa*. The virulotyping showed the presence of several relevant virulence genes such as T3SS (*eae*, *espA, espB, espD, espF, espI, espJ, and tir*) and other plasmid borne virulence relevant genes (*espP* and *exhA*) (Table 4), however a fully closed STEC genome was not recovered from any sample. While a fully closed STEC genome was not recovered, the detection of numerous virulence genes demonstrates centriflaken’s potential for comprehensive pathogen profiling, providing valuable insights for risk assessment in food safety contexts. AMR profiling showed the presence of a very diverse set of antimicrobial resistance genes per sample, from 2 to up to 20 genes, representing resistance to 10 antibiotic classes and multidrug efflux genes, underscores the potential of centriflaken for comprehensive pathogen characterization beyond just virulence factors (Supplementary Table 3).

**Table 4.** SSTEC virulotyping for 21 enriched agricultural waters by centriflaken.

| **Sample** | ***eae*** | ***saa*** | ***subA*** | ***exhA*** | ***espA*** | ***espB*** | ***espD*** | ***espF*** | ***espI*** | ***espJ*** | ***espP*** | ***tccP*** | ***tir*** |
| --- | --- | --- | --- | --- | --- | --- | --- | --- | --- | --- | --- | --- | --- |
| FAL00958 | + | - | - | + | + | + | + | - | - | + | - | - | + |
| FAL01198 | - | - | + | + | - | - | - | - | - | - | + | - | - |
| FAL01556 | - | - | - | + | - | - | - | - | - | - | - | - | - |
| FAL05306 | - | - | - | + | - | - | - | - | - | - | - | - | - |
| FAL11145 | - | - | - | + | - | - | - | - | - | - | - | - | - |
| FAL11151 | + | - | - | - | + | + | + | - | - | - | - | - | + |
| FAL11305 | - | - | - | + | - | - | - | - | - | - | - | - | - |
| FAL11351 | - | - | - | + | + | + | + | - | - | - | - | - | + |
| FAL11384 | - | - | - | + | - | - | - | - | - | - | - | - | - |
| FAL11387 | + | - | - | - | - | - | - | - | - | + | - | - | - |
| FAL11417 | - | - | - | + | - | - | - | - | - | - | - | - | - |
| FAL11425 | - | - | - | - | - | - | - | - | - | - | - | - | - |
| FAL16059 | - | - | - | + | - | - | - | - | - | - | - | - | - |
| FAL17459 | + | - | - | + | + | + | + | + | - | - | - | - | + |
| FAO71993 | - | - | - | + | - | - | - | - | - | - | - | - | - |
| FAO72169 | + | + | - | + | + | + | + | + | - | + | + | - | + |
| FAO96374 | - | - | - | + | - | - | - | - | - | - | - | - | - |
| FAO97359 | + | - | - | - | + | + | + | - | - | - | - | - | + |
| FAQ33890 | + | - | - | + | + | + | + | + | - | + | - | + | + |
| FAQ33923 | + | - | - | + | + | + | + | + | + | - | - | - | + |
| FAQ34034 | - | - | - | + | - | - | - | - | - | - | - | - | - |

## Discussion

### Importance of rapid and automated metagenomic analysis for food safety

Given the critical role of irrigation water in food production, rapid and accurate detection of STECs in these waters is essential but especially during outbreak investigations. Traditional detection methods, such as qPCR followed by extensive selective plating and whole genome sequencing (WGS), are labor-intensive and may take up to two weeks to confirm the correct identification of an isolate. Fast metagenomic analysis is crucial for accelerating this timeline. By integrating qPCR with long-read metagenomic sequencing of the pre-enrichment step, definitive detection and virulence characterization of STECs can be achieved within 3 to 4 days. While microbiological confirmation is still required, this approach potentially enables faster implementation of corrective actions since overall turnaround time is reduced by almost one week.

### centriflaken: addressing bioinformatic bottlenecks

However, the bioinformatic analysis of sequencing data [24] remains complex, involving many tools and steps to generate a complete STEC analysis. This process often requires up to 24 hours per sample -even when those samples analyses are run in parallel. To address this bottleneck, a workflow named centriflaken was developed to automate the entire process and to reduce the analysis time. centriflaken enables taxa-specific precision metagenomic analysis, *in silico* virulotyping, serotyping, and AMR prediction for STECs in a single, reproducible workflow. centriflaken consolidates the different steps of a precision metagenomic approach into a scalable, highly reproducible, and easy-to-install workflow using Nextflow [36]. It outputs final reports, summary plots and tables through a MultiQC HTML report for user friendly review as part of the pipeline. By automating and streamlining the analysis process, centriflaken not only saves time but also reduces the potential for human error, enhancing the reliability and reproducibility of STEC detection in complex environmental samples.

centriflaken uses many tools to provide a comprehensive analysis of ONT metagenomics samples. The associated tools were chosen based on the steps used for the analysis of ONT samples previously published [24] and by benchmark performance, *e.g.*, taxonomic classification and binning [37], *E. coli* reads extraction [24], assembly [33], taxonomic classification [34], and final *in silico* profiling [14, 38]. Built with a modular Nextflow DSL2 design, centriflaken is easily updated to include improved tools in future revisions or new ONT chemistries,such as the recent shift from R9.4.1 to R10.4.1.

### Performance benchmarking

centriflaken was first evaluated using previous analyzed data under Bioproject PRJNA639799 [24], and two ZymoBIOMICS microbial community datasets. For STEC-spiked samples, centriflaken achieved 100% genome completeness at concentrations >10⁷ CFU/ml and 85% at 10⁶ CFU/ml—mirroring the results from manual, stepwise analyses [24]. Serotyping, virulence gene detection, and AMR profiling were also consistent with previously reported outputs. In the ZymoBIOMICS microbial community dataset, centriflaken accurately identified the *E. coli* strain present in the community as serotype O33:H32 and detected four AMR-related genes, including *blaEC*, *mdtM*, *emrD*, and *acrF* [42]. In contrast, the *E. coli* genome was not recovered from the ZymoBIOMICS Microbial Community Standard II (Log Distribution), as its concentration was below the ∼10⁶ CFU threshold necessary for effective assembly [24].

In spiked agricultural water samples, centriflaken recovered STEC genomes with ≥85% completeness at ≥10⁶ CFU/ml (Table 1 and Figure 2). Although manual analysis produced slightly more contigs overall [24], the number of contigs corresponding to the STEC genome was equivalent, confirming the equivalency of both analysis workflows (Table 1). centriflaken also includes a step that extracts only the contigs belonging to the target taxa, streamlining the process and improving accuracy (Figure 1). The virulence profiles identified were consistent with prior results from Maguire et al. (2021) [24], confirming the accuracy and reproducibility of the workflow. (Table 2, Supplementary MultiQC Report 1),

### Real-world application and limitations

The performance of the centriflaken workflow was further evaluated using 21 real-life enriched agricultural water samples. The results showed that each sample contained multiple *E. coli* strains—ranging from 2 to 13—based on predicted serotype profiles. The presence of so many strains per sample, combined with their high concentrations as measured by qPCR (∼10⁶ CFU/ml), made it difficult to generate fully closed MAGs.

This finding highlights a key limitation of the methodology: recovering and characterizing STEC genomes *in silico* is challenging when samples are highly complex and contain multiple *E. coli* strains, especially when these strains outcompete the STEC target. Despite this challenge, three samples were identified as containing *stx* genes and adhesion genes such as *saa* or *eae*. An additional eight samples produced *E. coli* MAGs that harbored *eae* genes. The eae subtypes identified included: gamma-2 (in two samples), epsilon-7/Xi (in four samples), and beta-1 (in one sample). Regarding AMR prevalence in those 21 samples, between 2 and 20 genes were detected per sample. These genes potentially confer resistance to 10 antibiotic classes, including multidrug efflux systems. While these limitations highlight areas for future improvement, they also underscore the complexity of real-world samples and the importance of centriflaken’s ability to detect multiple strains simultaneously.

### Future directions and potential impact

centriflaken continues to evolve. The modular structure of the workflow setup of centriflaken using Nextflow allows further refinements as soon as future developments in ONT sequencing or metagenomics arise, as observed recently with the introduction of newer chemistries (R10.4.1) [43–51]. Nextflow-based parallelization enhances computational efficiency. With R10.4.1’s improved basecalling accuracy [43–48], there is growing potential to recover complete STEC genomes directly from complex samples— enabling phylogenetic comparisons to public genomes via the NCBI Pathogen Detection database. (https://www.ncbi.nlm.nih.gov/pathogens/).

ONT long reads further facilitate genome closure by resolving repetitive regions, as previously demonstrated with STEC EDL933 O157:H7 [24]. This is fully leveraged in centriflaken’s reassembly step, which focuses analysis on user-defined taxa to reduce computational load and maximize accuracy by only using the addressed taxa through a precision metagenomic approach as previously described [24]. centriflaken is publicly available at: https://github.com/CFSAN-Biostatstics/centriflaken.

centriflaken supports any user-defined taxon, including *Listeria monocytogenes* and *Salmonella*, expanding its utility for foodborne pathogen surveillance. All software dependencies are containerized for portability and reproducibility. The development of centriflaken as a useful tool in combination with metagenomic analysis is an important step in evaluation of a variety of samples associated with fresh produce. As a result, metagenomic analysis can now be completed within hours rather than weeks—providing precise detection and characterization of STEC and other pathogens in tested samples. While initially focused on STEC, centriflaken’s flexible design allows for adaptation to other foodborne pathogens. This versatility positions it as a valuable tool for comprehensive food safety monitoring, potentially revolutionizing how we approach pathogen detection and outbreak investigation.

### Looking to the future

centriflaken is currently available to Human Foods Program users via the HFP HPC cluster and GalaxyTrakr: https://www.protocols.io/view/centriflaken-an-automated-data-analysis-pipeline-f-kxygxzdbwv8j/v5. As ONT long-read metagenomics continues to mature, tools like centriflaken could play a transformative role in culture-independent pathogen detection and source tracking. Its portability makes it suitable for point-of-care testing and field deployment in food and water testing scenarios. Future studies should focus on optimizing centriflaken for even lower bacterial concentrations, integrating it with other emerging technologies like machine learning for predictive analytics, and validating its performance across a wider range of food and environmental matrices.

### Conclusion

centriflaken significantly accelerates and streamlines STEC detection in environmental samples. It provides a scalable, automated alternative to labor-intensive traditional workflows, delivering fast, reproducible, and accurate results. With continued enhancements and integration of newer ONT chemistries (e.g. R10.4.1), centriflaken may support real-time surveillance of foodborne pathogens directly from complex samples and shorten response times in outbreak scenarios. Furthermore, ongoing improvements and adaptations will potentially enhance its utility for food safety monitoring. In conclusion, centriflaken represents a substantial advancement in metagenomic analysis for food safety applications. By dramatically reducing analysis time and complexity while maintaining high accuracy, it has the potential to transform STEC detection and characterization, ultimately contributing to more rapid and effective food safety interventions.

## ACKNOWLEDGEMENTS

The study was supported by funding from the Chief Scientist-Challenge Grants Proposal #2021-1464 and the FDA Foods Program Intramural Funds.

## References

1. Taylor EV, Nguyen TA, Machesky KD, Koch E, Sotir MJ, Bohm SR, et al. Multistate outbreak of *Escherichia coli* O145 infections associated with romaine lettuce consumption, 2010. J Food Prot. 2013;76(6):939–44. doi: 10.4315/0362-028X.JFP-12-503 [doi].

2. Gould LH, Demma L, Jones TF, Hurd S, Vugia DJ, Smith K, et al. Hemolytic uremic syndrome and death in persons with *Escherichia coli* O157:H7 infection, foodborne diseases active surveillance network sites, 2000-2006. Clin Infect Dis. 2009;49(10):1480–5. Epub 2009/10/16. doi: 10.1086/644621. PubMed PMID: 19827953.

3. Hussein HS, Sakuma T. Shiga toxin-producing *Escherichia coli*: pre- and postharvest control measures to ensure safety of dairy cattle products. J Food Prot. 2005;68(1):199–207. Epub 2005/02/05. PubMed PMID: 15690827.

4. Sivapalasingam S, Friedman CR, Cohen L, Tauxe RV. Fresh produce: a growing cause of outbreaks of foodborne illness in the United States, 1973 through 1997. J Food Prot. 2004;67(10):2342–53. Epub 2004/10/29. doi: 10.4315/0362-028x-67.10.2342. PubMed PMID: 15508656.

5. Kulasekara BR, Jacobs M, Zhou Y, Wu Z, Sims E, Saenphimmachak C, et al. Analysis of the genome of the *Escherichia coli* O157:H7 2006 spinach-associated outbreak isolate indicates candidate genes that may enhance virulence. Infect Immun. 2009;77(9):3713–21. doi: IAI.00198-09 [pii];10.1128/IAI.00198-09 [doi].

6. Rangel JM, Sparling PH, Crowe C, Griffin PM, Swerdlow DL. Epidemiology of *Escherichia coli* O157:H7 outbreaks, United States, 1982-2002. Emerg Infect Dis. 2005;11(4):603–9. PubMed PMID: 15829201.

7. Teunis P, Takumi K, Shinagawa K. Dose response for infection by *Escherichia coli* O157:H7 from outbreak data. Risk Anal. 2004;24(2):401–7. Epub 2004/04/14. doi: 10.1111/j.0272-4332.2004.00441.x RISK441 [pii]. PubMed PMID: 15078310.

8. P F, KA L, H K, TS W. Genotypic and phenotypic changes in the emergence of *Escherichia coli* O157:H7. J Infect Dis. 1998;177(6):1750–3.

9. Mead PS, Griffin PM. *Escherichia coli* O157:H7. Lancet (London, England). 1998;352(9135):1207-12. PubMed PMID: Medline:9777854.

10. Bell BP, Goldoft M, Griffin PM, Davis MA, Gordon DC, Tarr PI, et al. A multistate outbreak of *Escherichia coli* O157:H7-associated bloody diarrhea and hemolytic uremic syndrome from hamburgers. The Washington experience. JAMA. 1994;272(17):1349–53.

11. Melton-Celsa AR. Shiga Toxin (Stx) Classification, Structure, and Function. Microbiol Spectr. 2014;2(4):EHEC-0024-2013. doi: 10.1128/microbiolspec.EHEC-0024-2013. PubMed PMID: 25530917; PubMed Central PMCID: PMCPMC4270005.

12. Ludwig K, Sarkim V, Bitzan M, Karmali MA, Bobrowski C, Ruder H, et al. Shiga toxin-producing *Escherichia coli* infection and antibodies against Stx2 and Stx1 in household contacts of children with enteropathic hemolytic-uremic syndrome. J Clin Microbiol. 2002;40(5):1773–82. doi: 10.1128/JCM.40.5.1773-1782.2002. PubMed PMID: 11980959; PubMed Central PMCID: PMCPMC130915.

13. Alhadlaq MA, Aljurayyad OI, Almansour A, Al-Akeel SI, Alzahrani KO, Alsalman SA, et al. Overview of pathogenic *Escherichia coli*, with a focus on Shiga toxin-producing serotypes, global outbreaks (1982-2024) and food safety criteria. Gut Pathog. 2024;16(1):57. Epub 20241007. doi: 10.1186/s13099-024-00641-9. PubMed PMID: 39370525; PubMed Central PMCID: PMCPMC11457481.

14. Joensen KG, Tetzschner AM, Iguchi A, Aarestrup FM, Scheutz F. Rapid and Easy In Silico Serotyping of *Escherichia coli* Isolates by Use of Whole-Genome Sequencing Data. J Clin Microbiol. 2015;53(8):2410–26. doi: JCM.00008-15 [pii];10.1128/JCM.00008-15 [doi].

15. Lorenz SC, Gonzalez-Escalona N, Kotewicz ML, Fischer M, Kase JA. Genome sequencing and comparative genomics of enterohemorrhagic *Escherichia coli* O145:H25 and O145:H28 reveal distinct evolutionary paths and marked variations in traits associated with virulence & colonization. BMC Microbiol. 2017;17(1):183. Epub 2017/08/24. doi: 10.1186/s12866-017-1094-3. PubMed PMID: 28830351; PubMed Central PMCID: PMCPMC5567499.

16. Mellmann A, Harmsen D, Cummings CA, Zentz EB, Leopold SR, Rico A, et al. Prospective genomic characterization of the German enterohemorrhagic *Escherichia coli* O104:H4 outbreak by rapid next generation sequencing technology. PLoS One. 2011;6(7):e22751. doi: 10.1371/journal.pone.0022751;PONE-D-11-11826.

17. Hedican EB, Medus C, Besser JM, Juni BA, Koziol B, Taylor C, et al. Characteristics of O157 versus non-O157 Shiga toxin-producing *Escherichia coli* infections in Minnesota, 2000-2006. Clin Infect Dis. 2009;49(3):358–64. Epub 2009/06/25. doi: 10.1086/600302. PubMed PMID: 19548834.

18. Bonnet R, Souweine B, Gauthier G, Rich C, Livrelli V, Sirot J, et al. Non-O157:H7 Stx2-producing *Escherichia coli* strains associated with sporadic cases of hemolytic-uremic syndrome in adults. J Clin Microbiol. 1998;36(6):1777–80.

19. Ogura Y, Ooka T, Iguchi A, Toh H, Asadulghani M, Oshima K, et al. Comparative genomics reveal the mechanism of the parallel evolution of O157 and non-O157 enterohemorrhagic *Escherichia coli*. Proc Natl Acad Sci U S A. 2009;106(42):17939–44. Epub 2009/10/10. doi: 0903585106 [pii] 10.1073/pnas.0903585106. PubMed PMID: 19815525; PubMed Central PMCID: PMC2764950.

20. Bielaszewska M, Prager R, Kock R, Mellmann A, Zhang W, Tschape H, et al. Shiga toxin gene loss and transfer in vitro and in vivo during enterohemorrhagic *Escherichia coli* O26 infection in humans. Appl Environ Microbiol. 2007;73(10):3144–50. Epub 2007/04/03. doi: AEM.02937-06 [pii] 10.1128/AEM.02937-06. PubMed PMID: 17400784; PubMed Central PMCID: PMC1907125.

21. Ogura Y, Ooka T, Asadulghani, Terajima J, Nougayrede JP, Kurokawa K, et al. Extensive genomic diversity and selective conservation of virulence-determinants in enterohemorrhagic *Escherichia coli* strains of O157 and non-O157 serotypes. Genome Biol. 2007;8(7):R138. Epub 2007/08/23. doi: gb-2007-8-7-r138 [pii] 10.1186/gb-2007-8-7-r138. PubMed PMID: 17711596; PubMed Central PMCID: PMC2323221.

22. Gonzalez-Escalona N, Kase JA. Virulence gene profiles and phylogeny of Shiga toxin-positive *Escherichia coli* strains isolated from FDA regulated foods during 2010-2017. PLoS One. 2019;14(4):e0214620. Epub 2019/04/02. doi: 10.1371/journal.pone.0214620. PubMed PMID: 30934002; PubMed Central PMCID: PMCPMC6443163.

23. Franz E, Delaquis P, Morabito S, Beutin L, Gobius K, Rasko DA, et al. Exploiting the explosion of information associated with whole genome sequencing to tackle Shiga toxin-producing *Escherichia coli* (STEC) in global food production systems. Int J Food Microbiol. 2014;187:57–72. doi: S0168-1605(14)00332-8;10.1016/j.ijfoodmicro.2014.07.002.

24. Maguire M, Kase JA, Roberson D, Muruvanda T, Brown EW, Allard M, et al. Precision long-read metagenomics sequencing for food safety by detection and assembly of Shiga toxin-producing *Escherichia coli* in irrigation water. PLoS One. 2021;16(1):e0245172. Epub 2021/01/15. doi: 10.1371/journal.pone.0245172. PubMed PMID: 33444384; PubMed Central PMCID: PMCPMC7808635.

25. Maguire M, Kase JA, Brown EWW, Allard MWW, Musser SMM, Gonzalez-Escalona N. Metagenomic survey of agricultural water using long read sequencing: Considerations for a successful analysis. Front Env Sci-Switz. 2022;10. doi: ARTN 830300 10.3389/fenvs.2022.830300. PubMed PMID: WOS:000844045400001.

26. Steele M, Odumeru J. Irrigation water as source of foodborne pathogens on fruit and vegetables. J Food Prot. 2004;67(12):2839–49. Epub 2005/01/07. doi: 10.4315/0362-028x-67.12.2839. PubMed PMID: 15633699.

27. Uyttendaele M, Jaykus L-A, Amoah P, Chiodini A, Cunliffe D, Jacxsens L, et al. Microbial hazards in irrigation water: standards, norms, and testing to manage use of water in fresh produce primary production. Comprehensive Reviews in Food Science and Food Safety. 2015;14(4):336–56. doi: 10.1111/1541-4337.12133.

28. Monaghan JM, Hutchison ML. Distribution and decline of human pathogenic bacteria in soil after application in irrigation water and the potential for soil-splash-mediated dispersal onto fresh produce. J Appl Microbiol. 2012;112(5):1007–19. Epub 2012/03/01. doi: 10.1111/j.1365-2672.2012.05269.x. PubMed PMID: 22372934.

29. Oliveira M, Viñas I, Usall J, Anguera M, Abadias M. Presence and survival of *Escherichia coli* O157:H7 on lettuce leaves and in soil treated with contaminated compost and irrigation water. Int J Food Microbiol. 2012;156(2):133–40. Epub 2012/04/10. doi: 10.1016/j.ijfoodmicro.2012.03.014. PubMed PMID: 22483400.

30. Allende A, Monaghan J. Irrigation water quality for leafy crops: a perspective of risks and potential solutions. Int J Environ Res Public Health. 2015;12(7):7457–77. Epub 2015/07/08. doi: 10.3390/ijerph120707457. PubMed PMID: 26151764; PubMed Central PMCID: PMCPMC4515668.

31. McCroskey LM, Hatheway CL, Woodruff BA, Greenberg JA, Jurgenson P. Type F botulism due to neurotoxigenic *Clostridium baratii* from an unknown source in an adult. J Clin Microbiol. 1991;29(11):2618–20.

32. Feng PW, Weagant S. D.; Jinneman, K. BAM Chapter 4A: diarrheagenic Escherichia coli 2020. Available from: https://www.fda.gov/food/laboratory-methods-food/bam-chapter-4a-diarrheagenic-escherichia-coli.

33. Kolmogorov M, Yuan J, Lin Y, Pevzner PA. Assembly of long, error-prone reads using repeat graphs. Nat Biotechnol. 2019;37(5):540–6. Epub 2019/04/03. doi: 10.1038/s41587-019-0072-8. PubMed PMID: 30936562.

34. Wood DE, Lu J, Langmead B. Improved metagenomic analysis with Kraken 2. Genome Biology. 2019;20(1):257. doi: 10.1186/s13059-019-1891-0.

35. Van Damme R, Holzer M, Viehweger A, Muller B, Bongcam-Rudloff E, Brandt C. Metagenomics workflow for hybrid assembly, differential coverage binning, metatranscriptomics and pathway analysis (MUFFIN). PLoS Comput Biol. 2021;17(2):e1008716. Epub 20210209. doi: 10.1371/journal.pcbi.1008716. PubMed PMID: 33561126; PubMed Central PMCID: PMCPMC7899367.

36. Di Tommaso P, Chatzou M, Floden EW, Barja PP, Palumbo E, Notredame C. Nextflow enables reproducible computational workflows. Nat Biotechnol. 2017;35(4):316–9. doi: 10.1038/nbt.3820. PubMed PMID: 28398311.

37. Kim D, Song L, Breitwieser FP, Salzberg SL. Centrifuge: rapid and sensitive classification of metagenomic sequences. Genome Res. 2016;26(12):1721–9. Epub 2016/11/18. doi: 10.1101/gr.210641.116. PubMed PMID: 27852649; PubMed Central PMCID: PMCPMC5131823.

38. Feldgarden M, Brover V, Gonzalez-Escalona N, Frye JG, Haendiges J, Haft DH, et al. AMRFinderPlus and the Reference Gene Catalog facilitate examination of the genomic links among antimicrobial resistance, stress response, and virulence. Sci Rep. 2021;11(1):12728. Epub 2021/06/18. doi: 10.1038/s41598-021-91456-0. PubMed PMID: 34135355; PubMed Central PMCID: PMCPMC8208984.

39. Doster E, Lakin SM, Dean CJ, Wolfe C, Young JG, Boucher C, et al. MEGARes 2.0: a database for classification of antimicrobial drug, biocide and metal resistance determinants in metagenomic sequence data. Nucleic Acids Res. 2020;48(D1):D561–D9. doi: 10.1093/nar/gkz1010. PubMed PMID: 31722416; PubMed Central PMCID: PMCPMC7145535.

40. Bortolaia V, Kaas RS, Ruppe E, Roberts MC, Schwarz S, Cattoir V, et al. ResFinder 4.0 for predictions of phenotypes from genotypes. J Antimicrob Chemother. 2020;75(12):3491–500. Epub 2020/08/12. doi: 10.1093/jac/dkaa345. PubMed PMID: 32780112; PubMed Central PMCID: PMCPMC7662176.

41. Gupta SK, Padmanabhan BR, Diene SM, Lopez-Rojas R, Kempf M, Landraud L, et al. ARG-ANNOT, a new bioinformatic tool to discover antibiotic resistance genes in bacterial genomes. Antimicrob Agents Chemother. 2014;58(1):212–20. Epub 20131021. doi: 10.1128/AAC.01310-13. PubMed PMID: 24145532; PubMed Central PMCID: PMCPMC3910750.

42. Joddha HB, Mathakiya RA, Joshi KV, Khant RB, Golaviya AV, Hinsu AT, et al. Profiling of antimicrobial resistance genes and integron from *Escherichia coli* isolates using whole genome sequencing. Genes (Basel). 2023;14(6). Epub 20230601. doi: 10.3390/genes14061212. PubMed PMID: 37372392; PubMed Central PMCID: PMCPMC10298372.

43. Hoffmann M, Jang JH, Tallent SM, Gonzalez-Escalona N. Single Laboratory Evaluation of the Q20+ Nanopore sequencing kit for bacterial outbreak investigations. Int J Mol Sci. 2024;25(22). Epub 20241105. doi: 10.3390/ijms252211877. PubMed PMID: 39595947; PubMed Central PMCID: PMCPMC11594029.

44. Gonzalez-Escalona N, Kwon HJ, Chen Y. Nanopore sequencing allows recovery of high-quality completely closed genomes of all *Cronobacter* species from powdered infant formula overnight enrichments. Microorganisms. 2024;12(12). Epub 20241122. doi: 10.3390/microorganisms12122389. PubMed PMID: 39770592; PubMed Central PMCID: PMCPMC11678115.

45. Lerminiaux N, Fakharuddin K, Mulvey MR, Mataseje L. Do we still need Illumina sequencing data? Evaluating Oxford Nanopore Technologies R10.4.1 flow cells and the Rapid v14 library prep kit for Gram negative bacteria whole genome assemblies. Canadian Journal of Microbiology. 2024. doi: 10.1139/cjm-2023-0175. PubMed PMID: WOS:001190425900001.

46. Sanderson ND, Hopkins KMV, Colpus M, Parker M, Lipworth S, Crook D, et al. Evaluation of the accuracy of bacterial genome reconstruction with Oxford Nanopore R10.4.1 long-read-only sequencing. Microb Genom. 2024;10(5). Epub 2024/05/07. doi: 10.1099/mgen.0.001246. PubMed PMID: 38713194.

47. Ritchie G, Chorlton SD, Matic N, Bilawka J, Gowland L, Leung V, et al. WGS of a cluster of MDR *Shigella sonnei* utilizing Oxford Nanopore R10.4.1 long-read sequencing. J Antimicrob Chemother. 2024;79(1):55–60. Epub 2023/11/15. doi: 10.1093/jac/dkad346. PubMed PMID: 37965757.

48. Hong YP, Chen BH, Wang YW, Teng RH, Wei HL, Chiou CS. The usefulness of nanopore sequencing in whole-genome sequencing-based genotyping of *Listeria monocytogenes* and *Salmonella enterica* serovar Enteritidis. Microbiol Spectr. 2024;12(7):e0050924. Epub 2024/05/29. doi: 10.1128/spectrum.00509-24. PubMed PMID: 38809017; PubMed Central PMCID: PMCPMC11218467.

49. Sanderson ND, Kapel N, Rodger G, Webster H, Lipworth S, Street TL, et al. Comparison of R9.4.1/Kit10 and R10/Kit12 Oxford Nanopore flowcells and chemistries in bacterial genome reconstruction. Microb Genom. 2023;9(1). Epub 2023/02/08. doi: 10.1099/mgen.0.000910. PubMed PMID: 36748454; PubMed Central PMCID: PMCPMC9973852.

50. Zhao W, Zeng W, Pang B, Luo M, Peng Y, Xu J, et al. Oxford nanopore long-read sequencing enables the generation of complete bacterial and plasmid genomes without short-read sequencing. Front Microbiol. 2023;14:1179966. Epub 2023/05/31. doi: 10.3389/fmicb.2023.1179966. PubMed PMID: 37256057; PubMed Central PMCID: PMCPMC10225699.

51. Sereika M, Kirkegaard RH, Karst SM, Michaelsen TY, Sorensen EA, Wollenberg RD, et al. Oxford Nanopore R10.4 long-read sequencing enables the generation of near-finished bacterial genomes from pure cultures and metagenomes without short-read or reference polishing. Nat Methods. 2022;19(7):823–6. Epub 2022/07/06. doi: 10.1038/s41592-022-01539-7. PubMed PMID: 35789207; PubMed Central PMCID: PMCPMC9262707 provides consulting and sequencing services. RHK, S.M.K. and T.Y.M. own shares in Oxford Nanopore Technologies PLC. The remaining author has no competing interests.

